# Computational Foundations of Natural Intelligence

**DOI:** 10.1101/166785

**Authors:** Marcel van Gerven

## Abstract

New developments in AI and neuroscience are revitalizing the quest to understanding natural intelligence, offering insight about how to equip machines with human-like capabilities. This paper reviews some of the computational principles relevant for understanding natural intelligence and, ultimately, achieving strong AI. After reviewing basic principles, a variety of computational modeling approaches is discussed. Subsequently, I concentrate on the use of artificial neural networks as a framework for modeling cognitive processes. This paper ends by outlining some of the challenges that remain to fulfill the promise of machines that show human-like intelligence.

## 1 Introduction

Understanding how mind emerges from matter is one of the great remaining questions in science. How is it possible that organized clumps of matter such as our own brains give rise to all of our beliefs, desires and intentions, ultimately allowing us to contemplate ourselves as well as the universe from which we originate? This question has occupied cognitive scientists who study the computational basis of the mind for decades. It also occupies other breeds of scientists. For example, ethologistis and psychologists focus on the complex behaviour exhibited by animals and humans whereas cognitive, computational and systems neuroscientists wish to understand the mechanistic basis of processes that give rise to such behaviour.

The ambition to understand natural intelligence as encountered in biological organisms can be contrasted with the motivation to build intelligent machines, which is the subject matter of artificial intelligence (AI). Wouldn’t it be amazing if we could build synthetic brains that are endowed with the same qualities as their biological cousins? This desire to mimick human-level intelligence by creating artificially intelligent machines has occupied mankind for many centuries. For instance, mechanical men and artificial beings appear in Greek mythology and realistic human automatons had already been developed in Hellenic Egypt (McCorduck, 2004). The engineering of machines that display human-level intelligence is also referred to as strong AI (Searle, 1980) or artificial general intelligence (AGI) (Uszkoreit et al., 2007), and was the original motivation that gave rise to the field of AI (Nilsson, 2005; Newell, 1991).

Excitingly, major advances in various fields of research now make it possible to attack the problem of understanding natural intelligence from multiple angles. From a theoretical point of view we have a solid understanding of the computational problems that are solved by our own brains (Dayan and Abbott, 2005). From an empirical point of view, technological breakthroughs allow us to probe and manipulate brain function in unprecedented ways, generating new neuroscientific insights about brain function (Chang, 2015). From an engineering perspective, we are finally able to build machines that learn to solve complex tasks, approximating and sometimes surpassing human-level performance (Jordan and Mitchell, 2015). Still, these efforts have not yet provided a full understanding of natural intelligence, nor did they give rise to machines whose reasoning capacity parallels the generality and flexibility of cognitive processing in biological organisms.

The core thesis of this paper is that natural intelligence can be better understood by the coming together of multiple complementary scientific disciplines (Gershman et al., 2015). This thesis is referred to as *the great convergence*. The advocated approach is to endow artificial agents with synthetic brains (i.e., cognitive architectures (Sun, 2004)) that mimick the thought processes that give rise to ethologically relevant behaviour in their biological counterparts. A motivation for this approach is given by Braitenberg’s law of uphill analysis and downhill invention, which states that it is much easier to understand a complex system by assembling it from the ground up, rather than by reverse engineering it from observational data (Braitenberg, 1986). These synthetic brains, which can be put to use in virtual or real-world environments, can then be validated against neuro-behavioural data and analysed using a multitude of theoretical tools. This approach not only elucidates our understanding of human brain function but also paves the way for the development of artificial agents that show truly intelligent behaviour.

The aim of this paper is to sketch the outline of a research program which marries the ambitions of neuroscientists to understand natural intelligence and AI researchers to achieve strong AI (Fig. 1). Before embarking on our quest to build synthetic brains as models of natural intelligence, we need to formalize what problems are solved by biological brains. That is, we first need to understand how adaptive behaviour ensues in animals and humans.

**Figure 1:**
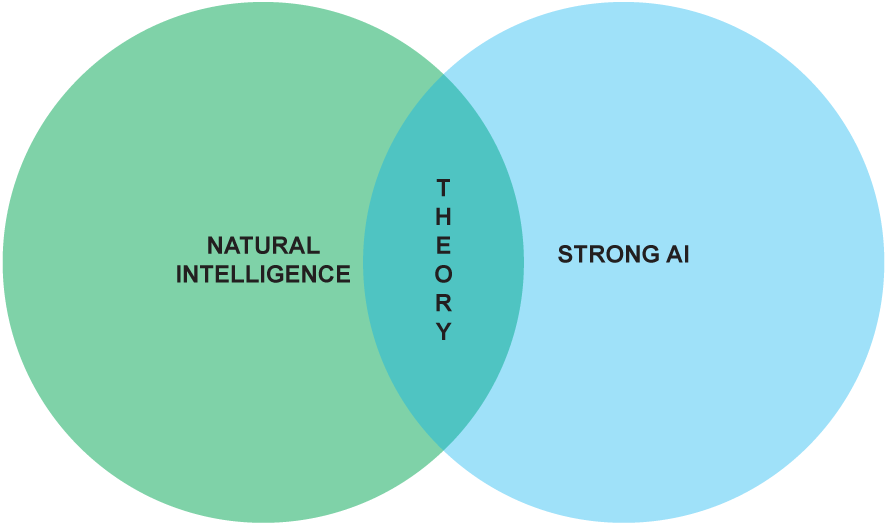
Understanding natural intelligence and achieving strong AI are seen as relying on the same theoretical foundations and require the convergence of multiple scientific and engineering disciplines.

## 2 Adaptive behaviour in biological agents

Ultimately, organisms owe their existence to the fact that they promote survival of their constituent genes; the basic physical and functional units of heredity that code for an organism (Dawkins, 2016). At evolutionary time scales, organisms developed a range of mechanisms which ensure that they live long enough such as to produce offspring. For example, single-celled protozoans already show rather complex ingestive, defensive and reproductive behavior, which is regulated by molecular signaling (Swanson, 2012; Sterling and Laughlin, 2016).

### 2.1 Why do we need a brain?

About 3.5 billion years ago, multicellular organisms started to appear. Multicellularity offers several competitive advantages over unicellularity. It allows organisms to increase in size without the limitations set by unicellularity and permits increased complexity by allowing cellular differentiation. It also increases life span since an organism can live beyond the demise of a single cell. At the same time, due to their increased size and complexity, multicellular organisms require more intricate mechanisms for signaling and regulation.

In multicellular organisms, behavior is regulated at multiple scales, ranging from intracellular molecular signaling all the way up to global regulation via the interactions between different organ systems. Hence, the nervous system allows for fast responses via electricochemical signaling and for slow responses by acting on the endocrine system. Nervous systems are found in almost all multicellular animals, but vary greatly in complexity. For example, the nervous system of the nematode roundworm Caenorhabditis elegans (C. elegans) is made up of 302 neurons and 7000 synaptic connections (White et al., 1986; Varshney et al., 2011). In contrast, the human brain contains about 20 billion neocortical neurons that are wired together via as many as 0.15 quadrillion synapses (Pakkenberg and Gundersen, 1997; Pakkenberg et al., 2003).

In vertebrates, the nervous system can be partitioned into the central nervous system (CNS), consisting of the brain and the spinal cord, and the peripheral nervous system (PNS), which connects the CNS to every other part of the body. The brain allows for centralised control and efficient information transmission. It can be partitioned into the forebrain, midbrain and hindbrain, each of which contain dedicated neural circuits that allow for integration of information and generation of coordinated activity. The spinal cord connects the brain to the body by allowing sensory and motor information to travel back and forth between the brain and the body. It also coordinates certain reflexes that bypass the brain altogether.

The interplay between the nervous system, the body and the environment is nicely captured by Swanson’s four system model of nervous system organization (Swanson, 2000), as shown in Figure 2. Briefly, the brain exerts centralized control on the body by sending commands to the motor system based on information received via the sensory system. It exerts this control by way of the cognitive system, which drives voluntary initiation of behavior, as well as the state system, which refers to the intrinsic activity that controls global behavioral state. The motor system can also be influenced directly by the sensory system via spinal cord reflexes. Output of the motor system induces visceral responses that affect bodily state as well as somatic responses that act on the environment. It is also able to drive the secretion of hormones that act more globally on the body. Both the body and the environment generate sensations that are processed by the sensory system. This closed-loop system, tightly coupling sensation, thought and action, is known as the *perception-action cycle* (Dewey, 1896; Sperry, 1952; Fuster, 2004).

**Figure 2:**
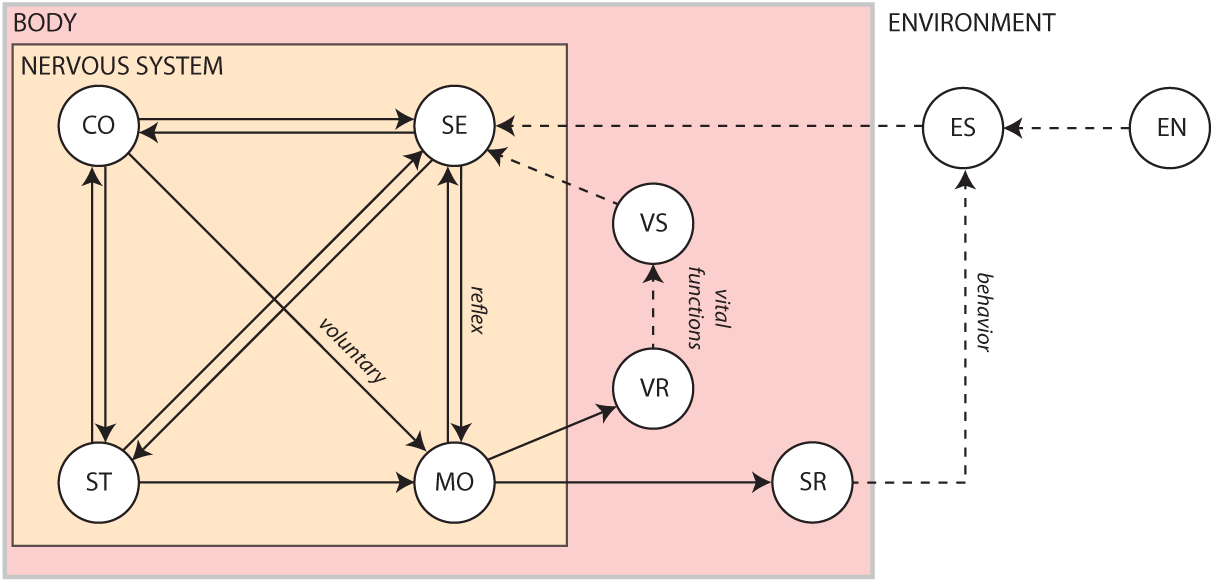
The four system model of nervous system organization. CO: Cognitive system; EN: Environment; ES: Environmental stimuli; MO; Motor system; SE: Sensory system; SR: Somatic responses; ST: Behavioural state system; VR: Visceral responses; VS: Visceral stimuli. Solid arrows show influences pertaining to the nervous system. Dashed arrows show interactions produced by the body or the environment (adapted from http://larrywswanson.com/?page_id=1523).

Summarizing, the brain, together with the spinal cord and the peripheral nervous system, can be seen as an organ that exploits sensory input such as to generate adaptive behavior through motor outputs. This ensures an organism’s long-term survival in a world that is dominated by uncertainty, as a result of partial observability, noise and stochasticity. The upshot of this interpretation is that action, which drives the generation of adaptive behaviour, is the ultimate reason why we have a brain in the first place. Citing Sperry (1952): “the entire output of our thinking machine consists of nothing but patterns of motor coordination.” To understand how adaptive behavior ensues, we therefore need to identify the ultimate causes that determine an agent’s actions (Tolman, 1932).

### 2.2 What makes us tick?

In biology, ultimately, all evolved traits must be connected to an organism’s survival. This implies that, from the standpoint of evolutionary psychology, natural selection favors those behaviors and thought processes that provide the organism with a selective advantage under ecological pressure (Barkow et al., 1992). Since causal links between behavior and long-term survival cannot be sensed or controlled directly, an agent needs to rely on other, directly accessible, ways to promote its survival. This can take the form of (1) evolving optimal sensors and effectors that allow it to maximize its control given finite resources and (2) evolving a behavioral repertoire that maximizes the information gained from the environment and generates optimal actions based on available sensory information.

In practice, behavior is the result of multiple competing needs that together provide an evolutionary advantage. These needs arise because they provide particular rewards to the organism. We distinguish *primary rewards*, *intrinsic rewards* and *extrinsic rewards*.

#### Primary rewards

Primary rewards are those necessary for the survival of one’s self and offspring, which includes homeostatic and reproductive rewards. Here, homeostasis refers to the maintenance of optimal settings of various biological parameters (e.g., regulation of temperature) (Cannon, 1929). A slightly more sophisticated concept is *allostasis*, which refers to the predictive regulation of biological parameters in order to prevent deviations rather than correcting them post hoc (Sterling, 2012). An organism can use its nervous system (muscle signaling) or endocrine system (endocrine signaling) to globally control or adjust the activities of many systems simultaneously. This allows for visceral responses that ensure proper functioning of an agent’s internal organs as well as basic drives such as ingestion, defense and reproduction that help ensure an agent’s survival (Tinbergen, 1951).

#### Intrinsic rewards

Intrinsic rewards are unconditioned rewards that are attractive and motivate behavior because they are inherently pleasurable (e.g., the experience of joy). The phenomenon of intrinsic motivation was first identified in studies of animals engaging in exploratory, playful and curiosity-driven behavior in the absence of external rewards or punishments (White, 1959).

#### Extrinsic rewards

Extrinsic rewards are conditioned rewards that motivate behavior but are not inherently pleasurable (e.g., praise or monetary reward). They acquire their value through learned association with intrinsic rewards. Hence, extrinsic motivation refers to our tendency to perform activities for known external rewards, whether they be tangible or psychological in nature (Brown, 2007).

Summarizing, the continual competition between multiple drives and incentives that have adaptive value to the organism and are realized by dedicated neural circuits is what ultimately generates behavior (Davies et al., 2012). In humans, the evolutionary and cultural pressures that shaped our own intrinsic and extrinsic motivations have allowed us to reach great achievements, ranging from our mastery of the laws of nature to expressions of great beauty as encountered in the liberal arts (Harari, 2015). The question remains how we can gain an understanding of how our brains generate the rich behavioral repertoire that can be observed in nature.

## 3 Understanding natural intelligence

In a way, the recipe for understanding natural intelligence and achieving strong AI is simple. If we can construct synthetic brains that mimick the adaptive behaviour displayed by biological brains in all its splendour then our mission has succeeded. This entails equipping synthetic brains with the same special purpose computing machinery encountered in real brains, solving those problems an agent may be faced with. In practice, of course, this is easier said than done given the incomplete state of our knowledge and the daunting complexity of biological systems.

### 3.1 Levels of analysis

The neural circuits that make up the human brain can be seen as special-purpose devices that together guarantee the selection of (near-)optimal actions. David Marr in particular advocated the view that the nervous system should be understood as a collection of information processing systems that solve particular problems an organism is faced with (Marr, 1982). His work gave rise to the field of computational neuroscience and has been highly influential in shaping ideas about neural information processing (Willshaw et al., 2015). Marr and Poggio (1976) proposed that an understanding of information processing systems should take place at distinct levels of analysis, namely the *computational level*, which specifies what problem the system solves, the *algorithmic level*, which specifies how the system solves the problem, and the *implementational level*, which specifies how the system is physically realized.

A canonical example of a three-level analysis is prey localization in the barn owl (Grothe, 2003). At the computational level, the owl needs to use auditory information to localize its prey. At the algorithmic level, this can be implemented by circuits composed of delay lines and coincidence detectors that detect inter-aural time differences (Jeffress, 1948). At the implementational level, neurons in the nucleus laminaris have been shown to act as coincidence detectors (Carr and Konishi, 1990).

Marr’s levels of analysis sidestep one important point, namely how a system gains the ability to solve a computational problem in the first place. That is, it is also crucial to understand how an organism (or species as a whole) is able to learn and evolve the computations and representations that allow it to survive in the natural world (Poggio, 2012). Learning itself takes place at the level of the individual organism as well as of the species. In the individual, one can observe lasting changes in the brain throughout its lifetime, which is referred to as neural plasticity. At the species level, natural selection is responsible for evolving the mechanisms that are involved in neural plasticity (Poggio, 2012). As argued by Poggio, an understanding at the level of learning in the individual and the species is sufficiently powerful to solve a problem and can thereby act as an explanation of natural intelligence. To illustrate the relevance of this revised model, in the prey localization example it would be imperative to understand how owls are able to adapt to changes in their environment (Huo and Murray, 2009), as well as how owls were equipped with such machinery during evolution.

Sun et al. (2005), in contrast, propose an alternative organisation of levels of cognitive modeling. They distinguish sociological, psychological, componential and physiological levels. The sociological level refers to the collective behavior of agents, including interactions between agents as well as their environment. It stresses the importance of socio-cultural processes in shaping cognition. The psychological level covers individual behaviors, beliefs, concepts, and skills. The componential level describes inter-agent processes specified in terms of Marr’s computational and algorithmic levels. Finally, the physiological level describes the biological substrate which underlies the generation of adaptive behavior, corresponding to Marr’s implementational level. It can provide valuable input about important computations and plausible architectures at a higher level of abstraction.

Without committing to a definitive stance on levels of analysis, all described levels provide important complementary perspectives concerning the modeling and understanding of natural intelligence.

### 3.2 Modeling approaches

The previous section suggests that different approaches to understanding natural intelligence and developing cognitive architectures can be taken depending on the levels of analysis one considers. We briefly review a number of core approaches.

#### Artificial life

Artificial life is a broad area of research encompassing various different modeling strategies which all have in common that they aim to explain the emergence of life and, ultimately, cognition in a bottom-up manner (Steels, 1993; Bedau, 2003).

A canonical example of an artificial life system is the cellular automaton, first introduced by von Neumann (1966) as an approach to understand the fundamental properties of living systems. Cellular automata change cell states based on simple local rules. They have been shown to be capable of acting as universal Turing machines, thereby giving them the capacity to compute any fixed partial computable function (Wolfram, 2002).

Figure 3 shows two examples of cellular automata. The left panel shows Gosper’s gliding gun, which is a structure which was used in a proof that Conway’s Game of Life is Turing complete (Gardner, 2001). SmoothLife (Rafler, 2011), shown in the right panel, is a continuous-space extension of the Game of Life whose emerging structures bear some superficial resemblance to structures that can be observed in biology. In principle, by virtue of their universality, cellular automata offer the capacity to explain how self-replicating adaptive (i.e. autopoeietic (Maturana and Varela, 1980)) systems emerge from basic rules. This bottom-up approach is also taken by physicists who aim to explain life and, ultimately, cognition purely from thermodynamic principles (Grinstein and Linsker, 2007; Dewar, 2005, 2003; Wissner-Gross and Freer, 2013; Perunov et al., 2014; Fry, 2017).

**Figure 3:**
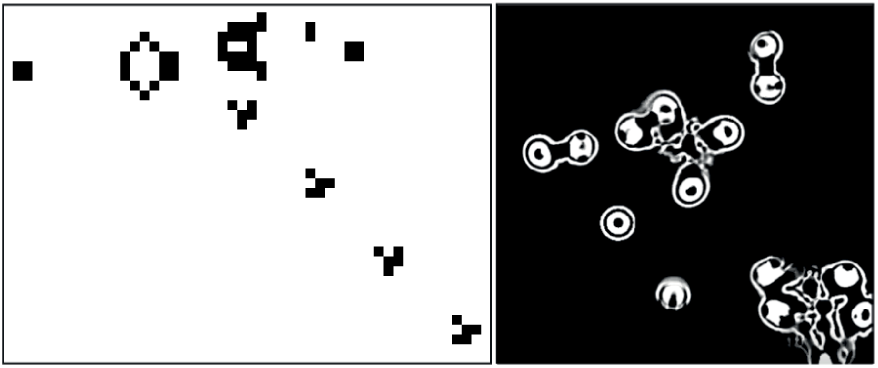
Examples of cellular automata. Left: Gosper’s glider gun in Conway’s Game of Life. Right: SmoothLife as a continuous-space extension of the Game of Life.

#### Biophysical modeling

A more direct way to model natural intelligence is to presuppose the existence of the building blocks of life which can be used to create realistic simulations of organisms in silico. The reasoning is that biophysically realistic models can eventually mimick the information processing capabilities of biological systems. An example thereof is the OpenWorm project which has as its ambition to understand how the behavior of C. elegans emerges from its underlying physiology purely via bottom-up biophysical modeling (Szigeti et al., 2014) (Fig. 4). It also acknowledges the importance of including not only a model of the worm’s nervous system but also of its body and environment in the simulation. That is, adaptive behavior depends on the organism being both embodied and embedded in the world (Anderson, 2003). If successful, then this project would constitute the first example of a digital organism.

**Figure 4:**
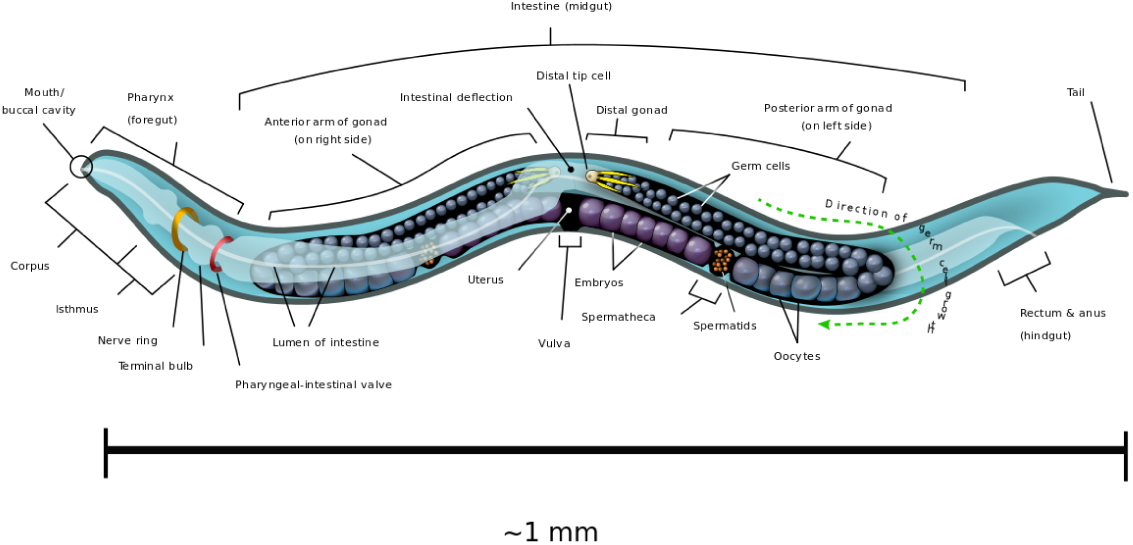
Body plan of C. elegans. The OpenWorm project aims to provide an accurate bottom-up simulation of the worm acting in its environment. Figure by K. D. Schroeder, CC BY-SA 3.0, https://commons.wikimedia.org/w/index.php?curid=26958836.

Its a long stretch from the worm’s 302 neurons to the 86 billion neurons that comprise the human brain (Herculano-Houzel and Lent, 2005). Still, researchers have set out to develop large-scale models of the human brain. Biophysical modeling can be used to create detailed models of neurons and their processes using coupled systems of differential equations. This strategy was used in the Blue Brain project (Markram, 2006) and its successor, the Human Brain Project (HBP) (Amunts et al., 2016). See de Garis et al. (2010) for a review of various artificial brain projects.

#### Connectionism

Connectionism refers to the explanation of cognition as arising from the interactions between simple (sub-symbolic) processing elements (Smolensky, 1987; Bechtel, 1993). It has close links to cybernetics, which focuses on the development of control structures from which intelligent behaviour emerges (Rid, 2016).

Connectionism came to be equated with the use of artificial neural networks that abstract away from the details of biological neural networks. An artificial neural network (ANN) is a computational model which is loosely inspired by the human brain as it consists of an interconnected network of simple processing units (artificial neurons) that learns from experience by modifying its connections. Alan Turing was one of the first to propose the construction of computing machinery out of trainable networks consisting of neuron-like elements (Copeland and Proudfoot, 1996). Marvin Minsky, one of the founding fathers of AI, is credited for building the first trainable ANN, called SNARC, out of tubes, motors, and clutches (Seising, 2017).

Artificial neurons can be considered abstractions of (populations of) neurons while the connections are taken to be abstractions of modifiable synaptic connections (Fig. 5). The behaviour of an artificial neuron is fully determined by the connection strengths as well as how input is transformed into output. Contrary to detailed biophysical models, ANNs make use of basic matrix operations and nonlinear transformations as their fundamental operations. In its most basic incarnation, an artificial neuron simply transforms its input x into a response *y* through an activation function *f*, as shown in Fig. 5. The activation function operates on an input activation which is typically taken to be the inner product between the input x and the parameters (weight vector) w of the artificial neuron. The weights are interpreted as synaptic strengths that determine how presynaptic input is translated into postsynaptic firing rate. This yields a simple linear-nonlinear mapping of the form

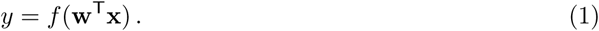

By connecting together multiple neurons, one obtains a neural network that implements some non-linear function y = f(x; *θ*), where the *f_i_* are nonlinear transformations and *θ* stands for the network parameters (i.e. weight vectors). After training a neural network, representations become encoded in a distributed manner as a pattern which manifests itself across all its neurons (Hinton et al., 1986).

**Figure 5:**
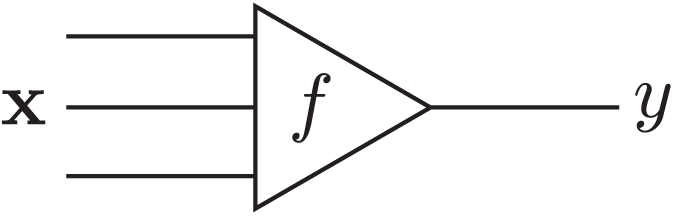
An artificial neuron receiving inputs x and generating output *y*.

Throughout the course of their history ANNs have fallen in and out of favor multiple times. At the same time, each next generation of neural networks has yielded new insights about how complex behaviour may emerge through the collective action of simple processing elements. Modern neural networks perform so well on several benchmark problems that they obliterate all competition in, e.g., object recognition (Krizhevsky et al., 2012), natural language processing (Sutskever et al., 2014), game playing (Mnih et al., 2015; Silver et al., 2016) and robotics (Levine et al., 2015), often matching and sometimes surpassing human-level performance (LeCun et al., 2015). Their success relies on combining classical ideas (Widrow and Lehr, 1990; Hochreiter and Schmidhuber, 1997; Lecun et al., 1998) with new algorithmic developments (Hinton et al., 2006; Srivastava et al., 2014; Ioffe and Szegedy, 2015; He et al., 2015), while using high-performance graphical processing units (GPUs) to massively speed up training of ANNs on big datasets (Raina et al., 2009).

#### Cognitivism

A completely different approach to the explanation of cognition as emerging from bottom-up principles is the view that cognition should be understood in terms of formal symbol manipulation. This computationalist view is associated with the cognitivist program which arose in response to earlier behaviorist theories. It embraces the notion that, in order to understand natural intelligence, one should study internal mental processes rather than just externally observable events. That is, cognitivism asserts that cognition should be defined in terms of formal symbol manipulation, where reasoning involves the manipulation of symbolic representations that refer to information about the world as acquired by perception.

This view is formalized by the physical symbol system hypothesis (Newell and Simon, 1976), which states that “a physical symbol system has the necessary and sufficient means for intelligent action.” This hypothesis implies that artificial agents, when equipped with the appropriate symbol manipulation algorithms, will be capable of displaying intelligent behavior. As Newell and Simon (1976) wrote, the physical symbol system hypothesis also implies that “the symbolic behavior of man arises because he has the characteristics of a physical symbol system.”

Cognitivism gave rise to cognitive science as well as artificial intelligence, and spawned various cognitive architectures such as ACT-R (Anderson et al., 2004) (see Fig. 6) and SOAR (Laird, 2012) that employ rule-based approaches in the search for a unified theory of cognition (Newell, 1991).^1^

**Figure 6:**
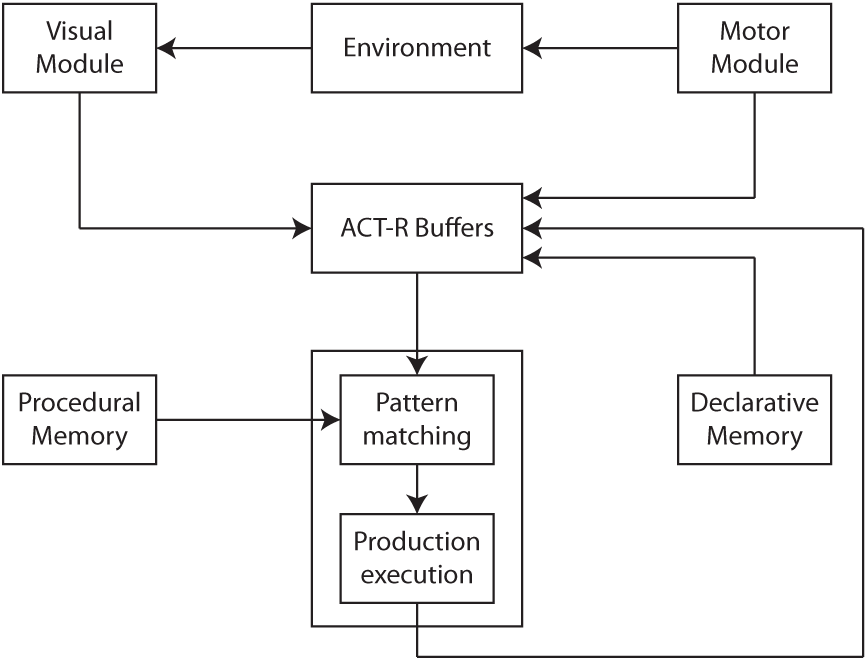
ACT-R as an example cognitive architecture which employs symbolic reasoning (adapted from http://act-r.psy.cmu.edu/about).

#### Probabilistic modeling

Modern cognitive science still embraces the cognitivist program but has since taken a probabilistic approach to the modeling of cognition. As stated by Griffiths et al. (2010), this probabilistic approach starts from the notion that the challenges faced by the mind are often of an inductive nature, where the observed data are not sufficient to unambiguously identify the process that generated them. This precludes the use of approaches that are founded on mathematical logic and requires a quantification of the state of the world in terms of degrees of belief as afforded by probability theory (Jaynes, 1988). The probabilistic approach operates by identifying a hypothesis space representing solutions to the inductive problem. It then prescribes how an agent should revise her belief in the hypotheses given the information provided by observed data. Hypotheses are typically formulated in terms of probabilistic graphical models that capture the independence structure between random variables of interest (Koller and Friedman, 2009). An example of such a graphical model is shown in Fig. 7.

**Figure 7:**
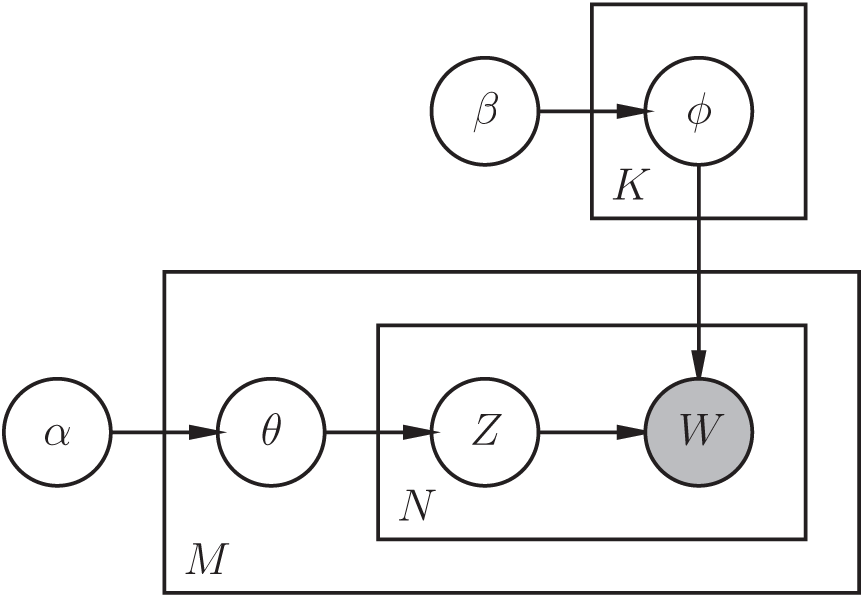
Example of a probabilistic graphical model capturing the statistical relations between random variables of interest. This particular model describes a smoothed version of latent Dirichlet allocation (Blei et al., 2003).

Belief updating in the probabilistic sense is realized by solving a statistical inference problem. Consider a set of of hypotheses ℋ that might explain the observed data. Let *p*(*h*) denote our belief in a hypothesis *h* ∈ ℋ, reflecting the state of the world, before observing any data (known as the *prior*). Let *p*(x | *h*) indicate the probability of observing data x if *h* were true (known as the *likelihood*). Bayes’ rule tells us how to update our belief in a hypothesis after observing data.

It states that the *posterior probability p*(*h* | x) assigned to *h* after observing x should be

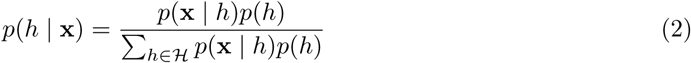

where the denominator is a normalizing constant known as the *evidence* or *marginal likelihood*.^2^ Importantly, it can be shown that degrees of belief are coherent only if they satisfy the axioms of probability theory (Ramsey, 1926).

The beauty of the probabilistic approach lies in its generality. It not only explains how our moment-to-moment percepts change as a function of our prior beliefs and incoming sensory data (Yuille and Kersten, 2006) but also places learning, as the construction of internal models, under the same umbrella by viewing it as an inference problem (MacKay, 2003). In the probabilistic framework, mental processes are modeled using algorithms for approximating the posterior (Koller and Friedman, 2009) and neural processes are seen as mechanisms for implementing these algorithms (Gershman and Beck, 2016).

The probabilistic approach also provides a basis for making optimal decisions under uncertainty. This is realized by extending probability theory with decision theory. According to decision theory, a rational agent ought to select that action which maximizes the expected utility (von Neumann and Morgenstern, 1953). This is known as the maximum expected utility (MEU) principle. In real-life situations, biological (and artificial) agents need to operate under bounded resources, trading off precision for speed and effort when trying to attain their objectives (Gigerenzer and Goldstein, 1996). This implies that MEU calculations may be intractable. Intractability issues have led to the development of algorithms that maximize a more general form of expected utility which incorporates the costs of computation. These algorithms can in turn be adapted so as to select the best approximation strategy in a given situation (Gershman et al., 2015). Hence, at the algorithmic level, it has been postulated that brains use approximate inference algorithms (Andrieu et al., 2003; Blei et al., 2016) such as to produce good enough solutions for fast and frugal decision making.

Summarizing, by appealing to Bayesian statistics and decision theory, while acknowledging the constraints biological agents are faced with, cognitive science arrives at a theory of bounded rationality that agents should adhere to. Importantly, this normative view dictates that organisms must operate as Bayesian inference machines that aim to maximize expected utility. If they do not, then, under weak assumptions, they will perform suboptimally. This would be detrimental from an evolutionary point of view.

### 3.3 Bottom-up emergence versus top-down abstraction

The aforementioned modeling strategies each provide an alternative approach towards understanding natural intelligence and achieving strong AI. The question arises which of these strategies will be most effective in the long run.

While the strictly bottom-up approach used in artificial life research may lead to fundamental insights about the nature of self-replication and adaptability, in practice it remains an open question how emergent properties that derive from a basic set of rules can reach the same level of organisation and complexity as can be found in biological organisms. Furthermore, running such simulations would be extremely costly from a computational point of view.

The same problem presents itself when using detailed biophysical models. That is, bottom-up approaches must either restrict model complexity or run simulations for limited periods of time in order to remain tractable (O’Reilly et al., 2012). Biophysical models additionally suffer from a lack of data. For example, the original aim of the Human Brain Project was to model the human brain within a decade (Markram et al., 2011). This ambition has been questioned since not enough bottom-up data may be available to estimate model parameters and the resulting models may fail to elucidate cognitive function. Izhikevich, reflecting on his simulation of another large biophysically realistic brain model (Izhikevich and Edelman, 2008), states: “Indeed, no significant contribution to neuroscience could be made by simulating one second of a model, even if it has the size of the human brain. However, I learned what it takes to simulate such a large-scale system.”^3^

Connectionist models, in contrast, abstract away from biophysical details, thereby making it possible to train large-scale models on large amounts of sensory data, allowing cognitively challenging tasks to be solved. Due to their computational simplicity, they are also more amenable to theoretical analysis (Hertz et al., 1991; Bishop, 1995). At the same time, connectionist models have been criticized for their inability to capture symbolic reasoning, their limitations when modeling particular cognitive phenomena, and their abstract nature, which restricts their biological plausibility (Dawson and Shamanski, 1994).

Cognitivism has been pivotal in the development of intelligent systems. However, it has also been criticized using the argument that systems which operate via formal symbol manipulation lack intentionality (Searle, 1980).^4^ Moreover, the representational framework that is used is typically constructed by a human designer. While this facilitates model interpretation, at the same time, this programmer-dependence may bias the system, leading to suboptimal solutions. That is, idealized descriptions may induce a semantic gap between perception and possible interpretation (Vernon et al., 2007).

The probabilistic approach to cognition is important given its ability to define normative theories at the computational level. At the same time, it has also been criticized for its treatment of cognition as if it is in the business of selecting some statistical model. Proponents of connectionism argue that computation-level explanations of behavior that ignore mechanisms associated with bottom-up emergence are likely to fall short (McClelland et al., 2010).

The different approaches provide complementary insights into the nature of natural intelligence. Artificial life informs about fundamental bottom-up principles, biophysical models make explicit how cognition is realized via specific mechanisms at the molecular and systems level, connectionist models show how problem solving capacities emerge from the interactions between basic processing elements, cognitivism emphasizes the importance of symbolic reasoning and probabilistic models inform how particular problems could be solved in an optimal manner.

Notwithstanding potential limitations, given their ability to solve complex cognitively challenging problems, connectionist models are taken to provide a promising starting point for understanding natural intelligence and achieving strong AI. They also naturally connect to the different modeling strategies. That is, they connect to artificial life principles by having network architectures emerge through evolutionary strategies (Salimans et al., 2017; Real et al., 2016) and connect to the biophysical level by viewing them as (rate-based) abstractions of biological neural networks (Dayan and Abbott, 2005). They also connect to the computational level by grounding symbolic representations in real-world sensory states (Harnad, 1990) and connect to the probabilistic approach through the observation that emergent computations effectively approximate Bayesian inference (Ambrogioni et al., 2017). It is for these reasons that, in the following, we will explore how ANNs, as canonical connectionist models, can be used to promote our understanding of natural intelligence.

## 4 ANN-based modeling of cognitive processes

We will now explore in more detail the ways in which ANNs can be used to understand and model aspects of natural intelligence. We start by addressing how neural networks can learn from data.

### 4.1 Learning

The capacity of brains to behave adaptively relies on their ability to modify their own behaviour based on changing circumstances. The appeal of neural networks stems from their ability to mimick this learning behaviour in an efficient manner by updating network parameters *θ* based on available data 𝒟 = {z^(1)^, …, z^(*N*)^}, allowing the construction of large models that are able to solve complex cognitive tasks.

Learning proceeds by making changes to the network parameters *θ* such that its output starts to agree more and more with the objectives of the agent at hand. This is formalized by assuming the existence of a *loss function* 𝒥 (*θ*) which measures the degree to which an agent deviates from its objectives. The loss 𝒥 is computed by running a neural network in forward mode (from input to output) and comparing the predicted output with the desired output. During its lifetime, the agent obtains data from its environment (i.e., sensations) by sampling from a data-generating distribution *p*_data_. The goal of an agent is to reduce the expected generalization error

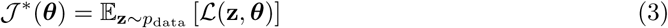

where ℒ is the incurred loss per datapoint z. In practice, an agent only has access to a finite number of datapoints which the agent experiences during its lifetime, yielding a training set 𝒟. This training set can be represented in the form of an empirical distribution *p̂*(**z**) which equals 1/*N* if **z** is equal to one of the *N* examples and zero otherwise. In practice, the aim therefore is to minimize the loss function

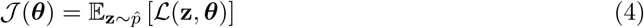

as an approximation of 𝒥* In reality, the brain is thought to optimize a multitude of loss functions pertaining to the many objectives it aims to achieve in concert (Marblestone et al., 2016).

Loss minimisation can be accomplished by making use of a gradient descent procedure. Let *θ* be the parameters of a neural network (i.e., the synaptic weights). We can define learning as a search for the optimal parameters *θ*\* based on available training data 𝒟 such that

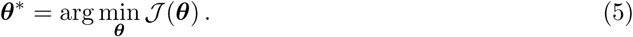

A convenient way to approximate *θ*\* is by measuring locally the change in slope of 𝒥 (*θ*) as a function of *θ* and taking a step in the direction of steepest descent. This procedure, known as *gradient descent*, is based on the observation that if 𝒥 is defined and differentiable in the neighbourhood of a point *θ* then 𝒥 decreases fastest if one goes from *θ* in the direction of the negative gradient −Δ_*θ*_ 𝒥 (*θ*). In other words, if we use the update rule

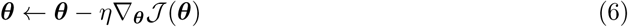

with small enough learning rate *η* then *θ* is guaranteed to converge to a (local) mimimum of 𝒥 (*θ*).^5^

Importantly, the gradient can be computed for arbitrary ANN architectures by running the network in backward mode (from output to input) and computing the gradient using automatic differentiation procedures. This forms the basis of the widely used backpropagation algorithm (Widrow and Lehr, 1990).

One might argue that the backpropagation algorithm fails to connect to learning in biology due to implausible assumptions such as the fact that forward and backward passes use the same set of synaptic weights. There are a number of responses here. First, one might hold the view that backpropagation is just an efficient way to obtain effective network architectures, without commiting to the biological plausibility of the learning algorithm per se. Second, if biologically plausible learning is the research objective then one is free to exploit other (Hebbian) learning schemes that may reflect biological learning more closely (Miconi, 2017). Finally, researchers have started to put forward arguments that backpropagation may not be that biologically implausible after all (Roelfsema and van Ooyen, 2005; Lillicrap et al., 2016; Scellier and Bengio, 2017).

### 4.2 Perceiving

One of the core skills any intelligent agent should possess is the ability to recognize patterns in its environment. The world around us consists of various objects that may carry significance. Being able to recognize edible food, places that provide shelter, and other agents will all aid survival.

Biological agents are faced with the problem that they need to be able to recognize objects from raw sensory input (vectors in ℝ^*n*^). How can a brain use the incident sensory input to learn to recognize those things that are of relevance to the organism? Recall the artificial neuron formulation *y* = *f*(w^T^x). By learning proper weights w, this neuron can learn to distinguish different object categories. This is essentially equivalent to a classical model known as the perceptron (Rosenblatt, 1958), which was used to solve simple pattern recognition problems via a simple error-correction mechanism. It also corresponds to a basic linear-nonlinear (LN) model which has been used extensively to model and estimate the receptive field of a neuron or a population of neurons (van Gerven, 2017).

Single-layer ANNs such as the perceptron are capable of solving interesting learning problems. At the same time, they are limited in scope since they can only solve linearly separable classification problems (Minsky and Papert, 1969). To overcome the limitations of the perceptron we can extend its capabilities by relaxing the constraint that the inputs are directly coupled to the outputs. A multilayer perceptron (MLP) is a feedforward network which generalizes the standard perceptron by having a hidden layer that resides between the input and the output layers. We can write an MLP with multiple output units as

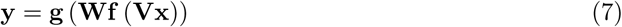

where **V** denotes the hidden layer weights and **W** denotes the output layer weights. By introducing a hidden layer, MLPs gain the ability to learn internal representations (Rumelhart et al., 1986). Importantly, an MLP can approximate any continuous function to an arbitrary degree of accuracy, given a sufficiently large but finite number of hidden neurons (Hornik, 1991; Cybenko, 1989).

Complex systems tend to be hierarchical and modular in nature (Simon, 1962). The nervous system itself can be thought of as a hierarchically organized system. This is exemplified by Felleman & van Essen’s hierarchical diagram of visual cortex (Felleman and Van Essen, 1991), the proposed hierarchical organisation of prefrontal cortex (Badre, 2008), the view of the motor system as a behavioral control column (Swanson, 2000) and the proposition that anterior and posterior cortex reflect hierarchically organised executive and perceptual systems (Fuster, 2001). Representations at the top of these hierarchies correspond to highly abstract statistical invariances that occupy our ecological niche (Barlow, 2009; Quian Quiroga et al., 2005). A hierarchy can be modeled by a deep neural network (DNN) composed of multiple hidden layers (LeCun et al., 2015), written as

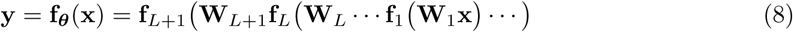

where **W**_*l*_ is the weight matrix associated with layer *l*. Even though an MLP can already approximate any function to an arbitrary degree of precision, it has been shown that many classes of functions can be represented much more compactly using thin and deep neural networks compared to shallow and wide neural networks (Bengio and Lecun, 2007; Delalleau and Bengio, 2011; Roux and Bengio, 2010; Bengio, 2009; Mhaskar et al., 2016).

A DNN corresponds to a stack of LN models, generalizing the concept of basic receptive field models. They have been shown to yield human-level performance on object categorisation tasks (Krizhevsky et al., 2012). The latest DNN incarnations are even able to predict the cognitive states of other agents. One example is the prediction of (apparent) personality traits from multimodal sensory input (Güçlütürk et al., 2016). Deep architectures have been used extensively in neuroscience to model hierarchical processing (Selfridge, 1959; Fukushima, 1980; Riesenhuber and Poggio, 1999; Fukushima, 2013; Lehky and Tanaka, 2016). Interestingly, it has been shown that the representations encoded in DNN layers correspond to the representations that are learned by areas that make up the sensory hierarchies of biological agents (Güçlü and van Gerven, 2015, 2017; Güçlü et al., 2016). Multiple reviews discuss this use of DNNs in sensory neuroscience (Cox and Dean, 2014; Kriegeskorte, 2015; Robinson and Rolls, 2015; van Gerven, 2017; Yamins and DiCarlo, 2016; Marblestone et al., 2016; Vanrullen, 2017; Peelen and Downing, 2017; Kietzmann et al., 2017).

### 4.3 Remembering

Being able to perceive the environment also implies that agents can store and retrieve past knowledge about objects and events in their surroundings. In the feedforward networks considered in the previous section, this knowledge is encoded in the synaptic weights as a result of learning. Memories of the past can also be stored, however, in moment-to-moment neural activity patterns. This does require the availability of lateral or feedback connections in order to enable recurrent processing (Singer, 2013; Maass, 2016). Recurrent processing can be implemented by a recurrent neural network (RNN) (Elman, 1990; Jordan, 1990), defined by

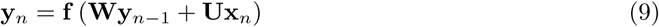

such that the neuronal activity at time *n* depends on the activity at time *n* – 1 as well as instantaneous bottom-up input. RNNs can be interpreted as numerical approximations of differential equations that describe rate-based neural models (Dayan and Abbott, 2005) and have been shown to be universal approximators of dynamical systems (Funahashi and Nakamura, 1993).^6^ Their parameters can be estimated using a variant of backpropagation, referred to as backpropagation through time (Mozer, 1989).

When considering perception, feedforward architectures may seem sufficient. For example, the onset latencies of neurons in monkey inferior-temporal cortex during visual processing are about 100 ms (Thorpe and Fabre-Thorpe, 2001), which means that there is ample time for the transmission of just a few spikes. This suggests that object recognition is largely an automatic feedforward process (Vanrullen, 2007). However, recurrent processing is important in perception as well since it provides the ability to maintain state. This is important in detecting salient features in space and time (Joukes et al., 2014), as well as for integrating evidence in noisy or ambiguous settings (O’Reilly et al., 2013). Moreover, perception is strongly influenced by top-down processes, as mediated by feedback connections (Gilbert and Li, 2013). RNNs have also been used to model working memory (Miconi, 2017) as well as hippocampal function, which is involved in a variety of memory-related processes (Willshaw et al., 2015; Kumaran et al., 2016).

A special kind of RNN is the Hopfield network (Hopfield, 1982), where **W** is symmetric and **U** = **0**. Learning in a Hopfield net is based on a Hebbian learning scheme. Hopfield nets are attractor networks that converge to a state that is a local minimum of an energy function. They have been used extensively as models of associative memory (Wills et al., 2005). It has even been postulated that dreaming can be seen as an unlearning process which gets rid of spurious minima in attractor networks, thereby improving their storage capacity (Crick and Mitchison, 1983).

### 4.4 Acting

As already described, the ability to generate appropriate actions is what ultimately drives behavior. In real-world settings, such actions typically need to be inferred from reward signals *r_t_* provided by the environment. This is the subject matter of reinforcement learning (RL) (Sutton and Barto, 1998). Define a policy π(*s, a*) as the probability of selecting an action *a* given a state *s*. Let the return 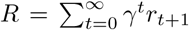 be the total reward accumulated in an episode, with *γ* a discount factor that downweighs future rewards. The goal in RL is to identify an optimal policy *π*\* that maximizes the expected return

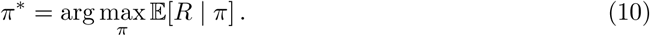

Reinforcement learning algorithms have been crucial in training neural networks that have the capacity to act. Such networks learn to generate suitable actions purely by observing the rewards entailed by previously generated actions. RL algorithms come in model-free and model-based variants. In the model-free setting, optimal actions are learned purely based on the reward that is gained by performing actions in the past. In the model-based setting, in contrast, an explicit model of the environment is used to predict the consequences of actions that are being executed. Importantly, model-free and model-based reinforcement learning approaches have clear correspondences with habitual and goal-directed learning in neuroscience (Daw, 2012; Buschman et al., 2014).

Various model-free reinforcement learning approaches have been used to develop a variety of neural networks for action generation. For example, Q-learning was used to train networks that play Atari games (Mnih et al., 2015) and policy gradient methods have been used to play board games (Silver et al., 2016) and solve problems in (simulated) robotics (Silver et al., 2014; Schulman et al., 2015), effectively closing the perception-action cycle. Evolutionary strategies are also proving to become an useful approach for solving challenging control problems (Salimans et al., 2017). Similar successes have been achieved using model-based reinforcement learning approaches (Schmidhuber, 2015; Mujika, 2016; Santana and Hotz, 2016).

Another important ingredient required for generating optimal actions is recurrent processing, as described in the previous section. Action generation must depend on the ability to integrate evidence over time since, otherwise, we are guaranteed to act suboptimally. That is, states that are qualitatively different can appear the same to the decision maker, leading to suboptimal policies. Consider for example the sensation of a looming object. The optimal decision depends crucially on whether this object is approaching or receding, which can only be determined by taking past sensations into account. This phenomenon is known as perceptual aliasing (Whitehead and Ballard, 1991).

A key ability of biological organisms which requires recurrent processing is their ability to navigate in their environment, as mediated by the hippocampal formation (Moser et al., 2015). Recent work shows that particular characteristics of hippocampal place cells, such as stable tuning curves that remap between environments, are recovered by training neural networks on navigation tasks (Kanitscheider and Fiete, 2016). The ability to integrate evidence also allows agents to selectively sample the environment, such as to maximise the amount of information gained. This process, known as active sensing, is crucial for understanding perceptual processing in biology (Yarbus, 1967; Regan and Noë, 2001; Schroeder et al., 2010; Friston et al., 2010; Schroeder et al., 2010; Gordon and Ahissar, 2012). Active sensing, in the form of saccade planning, has been implemented using a variety of recurrent neural network architectures (Larochelle and Hinton, 2010; Gregor et al., 2014; Mnih et al., 2014). RNNs that implement recurrent processing have also been used to model various other action-related processes such as timing (Laje and Buonomano, 2013), sequence generation (Rajan et al., 2015) and motor control (Sussillo et al., 2015).

Recurrent processing and reinforcement learning are also essential in modeling higher-level processes, such as cognitive control as mediated by frontal brain regions (Miller and Cohen, 2001; Fuster, 2001). Examples are models of context-dependent processing (Mante et al., 2013) and perceptual decision-making (Carnevale et al., 2015), In general, RNNs that have been trained using RL on a variety of cognitive tasks have been shown to yields properties that are consistent with phenomena observed in biological neural networks (Miconi, 2017; Song et al., 2016).

### 4.5 Predicting

Modern theories of human brain function appeal to the idea that the brain can be viewed as a prediction machine, which is in the business of continuously generating top-down predictions that are integrated with bottom-up sensory input (Yuille and Kersten, 2006; Lee and Mumford, 2003; Clark, 2013; Summerfield and de Lange, 2014). This view of the brain as a prediction machine that performs unconscious inference has a long history, going back to the seminal work of Alhazen and Helmholtz (Hatfield, 2002). Modern views cast this process in terms of Bayesian inference, where the brain is updating its internal model of the environment in order to explain away the data that impinge upon its senses, also referred to as the Bayesian brain hypothesis (Jaynes, 1988; Doya et al., 2006). The same reasoning underlies the free-energy principle, which assumes that biological systems minimise a free energy functional of their internal states that entail beliefs about hidden states in their environment (Friston, 2010). Predictions can be seen as central to the generation of adaptive behavior, since anticipating the future will allow an agent to select appropriate actions in the present (Moulton and Kosslyn, 2009; Schacter et al., 2007).

Prediction is central in model-based RL approaches since it requires agents to plan their actions by predicting the outcomes of future actions (Daw, 2012). This is strongly related to the notion of preplay of future events subserving path planning (Corneil and Gerstner, 2015). Such preplay has been observed in hippocampal place cell sequences (Dragoi and Tonegawa, 2011), giving further support to the idea that the hippocampal formation is involved in goal-directed navigation (Corneil and Gerstner, 2015). Prediction also allows an agent to prospectively act on expected deviations from optimal conditions. This focus on error-correction and stability is also prevalent in the work of the cybernetic movement (Ashby, 1952). Note further that predictive processing connects to the concept of allostasis, where the agent is actively trying to predict future states such as to minimize deviations from optimal homeostatic conditions. It is also central to optimal feedback control theory, which assumes that the motor system corrects only those deviations that interfere with task goals (Todorov and Jordan, 2002).

The notion of predictive processing has been very influential in neural network research. For example, it provides the basis for predictive coding models that introduce specific neural network architectures in which feedforward connections are used to transmit the prediction errors that result from discrepancies between top-down predictions and bottom-up sensations (Rao and Ballard, 1999; Huang and Rao, 2011). It also led to the development of a wide variety of generative models that are able to predict their sensory states, also referred to as fantasies (Hinton, 2013). Such fantasies may play a role in understanding cognitive processing involved in imagery, working memory and dreaming. In effect, these models aim to estimate a distribution over latent causes z in the environment that explain observed sensory data x. In this setting, the most probable explanation is given by

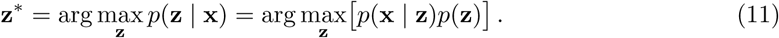

Generative models also offer a way to perform unsupervised learning, since if a neural network is able to generate predictions then the discrepancy between predicted and observed stimuli can serve as a teaching signal. A canonical example is the Boltzmann machine, which is a stochastic variant of a Hopfield network that is able to discover regularities in the training data using a simple unsupervised learning algorithm (Ackley et al., 1985; Hinton and Sejnowski, 1983). Another classical example is the Helmholtz machine, which incorporates both bottom-up and top-down processing (Dayan et al., 1995). Other, more recent examples of ANN-based generative models are deep belief networks (Hinton et al., 2006), variational autoencoders (Kingma and Welling, 2014) and generative adversarial networks (Goodfellow et al., 2014). Recent work has started to use these models to predict future sensory states from current observations (Mathieu et al., 2016; Lotter et al., 2016; Xue et al., 2016).

### 4.6 Reasoning

While ANNs are now able to solve complex tasks such as acting in natural environments or playing difficult board games, one could still argue that they are ‘just’ performing sophisticated pattern recognition rather than showing the symbolic reasoning abilities that characterize our own brains. The question of whether connectionist systems are capable of symbolic reasoning has a long history, and has been debated by various researchers in the cognitivist (symbolic) program (Pinker and Mehler, 1988). We will not settle this debate here but point out that efforts are underway to endow neural networks with sophisticated reasoning capabilities.

One example is the development of ‘differentiable computers’ that learn to implement algorithms based on a finite amount of training data (Graves et al., 2014; Weston et al., 2015; Vinyals et al., 2017). The resulting neural networks perform variable binding and are able to deal with variable length structures (Graves et al., 2014), which are two objections that were originally raised against using ANNs to explain cognitive processing (Fodor and Pylyshyn, 1988).

Another example is the development of neural networks that can answer arbitrary questions about text (Bordes et al., 2015), images (Agrawal et al., 2016) and movies (Tapaswi et al., 2015), thereby requiring deep semantic knowledge about the experienced stimuli. Recent models (Johnson et al., 2017) have also been shown to be capable of compositional reasoning (Lake et al., 2016), which is an important ingredient for explaining the systematic nature of human thought (Fodor and Pylyshyn, 1988). These architectures often make use of distributional semantics, where words are encoded as real vectors that capture word meaning (Mikolov et al., 2013; Ferrone and Zanzotto, 2017).

Several other properties characterize human thought processes, such as intuitive physics, intuitive psychology, relational reasoning and causal reasoning (Lake et al., 2016; Kemp and Tenenbaum, 2008). Another crucial hallmark of intelligent systems is that they are able to explain what they are doing (Brachman, 2002). This requires agents to have a deep understanding of their world.^7^ These properties should be replicated in neural networks if they are to serve as accurate models of natural intelligence. New neural network architectures are slowly starting to take steps in this direction (e.g., (Santoro et al., 2017; Zhu et al., 2017; Louizos et al., 2017)).

## 5 Towards strong AI

We have reviewed the computational foundations of natural intelligence and outlined how ANNs can be used to model a variety of cognitive processes. However, our current understanding of natural intelligence remains limited and strong AI has not yet been attained. In the following, we will touch upon a number of important topics that will be of importance for eventually reaching these goals.

### 5.1 Surviving in complex environments

Contemporary neural network architectures tend to excel at solving one particular problem well. However, in practice, we want to arrive at intelligent machines that are able to survive in complex environments. This requires the agent to deal with high-dimensional naturalistic input, be able to solve multiple tasks depending on context, and devise optimal strategies to ensure long-term survival.

The research community has embraced these desiderata by creating virtual worlds that allow development and testing of neural network architectures (e.g., (Beattie et al., 2016; Kempka et al., 2016; Todorov et al., 2012; Synnaeve et al., 2016; Brockman et al., 2016)).^8^ While most work in this area has focused on environments with fully observable states, reward functions with low delay, and small action sets, research is shifting towards environments that are partially observable, require long-term planning, show complex dynamics and have noisy and high-dimensional control interfaces (Synnaeve et al., 2016).

A particular challenge in these naturalistic environments is that networks need to be able to exhibit continual (life-long) learning (Thrun and Mitchell, 1995), adapting continuously to the current state of affairs. This is difficult due to the phenomenon of catastrophic forgetting (Mccloskey and Cohen, 1986; French, 1999), where previously acquired skills are overwritten by ongoing modification of synaptic weights. Recent algorithmic developments attenuate the detrimental effects of catastrophic forgetting (Kirkpatrick et al., 2015; Zenke et al., 2015), offering a (partial) solution to the stability versus plasticity dilemma (Abraham and Robins, 2005). Life-long learning is further complicated by the exploration-exploitation dilemma, where agents need to decide on whether to accrue either information or reward (Cohen et al., 2007). Another challenge is the fact that reinforcement learning of complex actions is notoriously slow. Here, progress is being made using networks that make use of differentiable memories (Pritzel et al., 2017; Santoro et al., 2016). Survival in complex environments also requires that agents learn to perform multiple tasks well. This learning process can be facilitated through multitask learning (Caruana, 1997) (also referred to as learning to learn (Baxter, 1998) or transfer learning (Pan and Fellow, 2009)), where learning of one task is facilitated by knowledge gained through learning to solve another task. Multitask learning has been shown to improve convergence speed and generalization to unseen data (Scholte et al., 2017). Finally, effective learning also calls for agents that can generalize to cases that were not encountered before, which is known as zero-shot learning (Palatucci et al., 2009), and can learn from rare events, which is known as one-shot learning (Fei-fei et al., 2006; Kaiser and Roy, 2017; Vinyals et al., 2016).

While the use of virtual worlds allows for testing the capabilities of artificial agents, it does not guarantee that the same agents are able to survive in the real world (Tobin et al., 2017). That is, there may exist a reality gap (Tobin et al., 2017), where skills acquired in virtual worlds do not carry over to the real world. In contrast to virtual worlds, acting in the real world requires the agent to deal with unforeseen circumstances resulting from the complex nature of reality, the agent’s need for a physical body, as well as its engagement with a myriad of other agents (Anderson, 2003). Moreover, the continuing interplay between an organism and its environment may itself shape and, ultimately, determine cognition (H. Maturana and F. Varela, 1987; Gibson, 1979; Brooks, 1996; Edelman, 2015). Effectively dealing with these complexities may not only require plasticity in individual agents but also the incorporation of developmental change, as well as learning at evolutionary time scales (Marcus, 2009). From a developmental perspective, networks can be more effectively trained by presenting them with a sequence of increasingly complex tasks, instead of immediately requiring the network to solve the most complex task (Elman, 1993). This process is known as curriculum learning (Bengio et al., 2009) and is analogous to how a child learns by decomposing problems into simpler subproblems (Turing, 1950). Evolutionary strategies have also been shown to be effective in learning to solve challenging control problems (Salimans et al., 2017). Finally, to learn about the world, we may also turn towards cultural learning, where agents can offload task complexity by learning from each other (Bengio, 2012).

As mentioned in Section 2.2, adaptive behavior is the result of multiple competing drives and motivations that provide primary, intrinsic and extrinsic rewards. Hence, one strategy for endowing machines with the capacity to survive in the real world is to equip neural networks with drives and motivations that ensure their long-term survival.^9^ In terms of primary rewards, one could conceivably provide artificial agents with the incentive to minimize computational resources or maximize offspring via evolutionary processes (Stanley and Miikkulainen, 2002; Floreano et al., 2008; Gauci and Stanley, 2010). In terms of intrinsic rewards, one can think of various ways to equip agents with the drive to explore the environment (Oudeyer, 2007). We briefly describe a number of principles that have been proposed in the literature. Artificial curiosity assumes that internal reward depends on how boring an environment is, with agents avoiding fully predictable and unpredictably random states (Schmidhuber, 2003, 1991; Pathak et al., 2017). A related notion is that of information-seeking agents (Bachman et al., 2016). The autotelic principle formalizes the concept of flow where an agent tries to maintain a state were learning is challenging, but not overwhelming (Csikszentmihalyi, 1975; Steels, 2004). The free-energy principle states that an agent seeks to minimize uncertainty by updating its internal model of the environment and selecting uncertainty-reducing actions (Friston, 2009, 2010). Empowerment is founded on information-theoretic principles and quantifies how much control an agent has over its environment, as well as its ability to sense this control (Klyubin et al., 2005a,b; Salge et al., 2013). In this setting, intrinsically motivated behavior is induced by the maximization of empowerment. Finally, various theories embrace the notion that optimal prediction of future states drives learning and behavior (Kaplan and Oudeyer, 2004; Der et al., 1999; Ay et al., 2008). In terms of extrinsic rewards, one can think of imitation learning, where a teacher signal is used to inform the agent about its desired outputs (Schaal, 1999; Duan et al., 2017).

### 5.2 Bridging the gap between artificial and biological neural networks

To reduce the gap between artificial and biological neural networks, it makes sense to assess their operation on similar tasks. This can be done either by comparing the models at a neurobiological level or at a behavioral level. The former refers to comparing the internal structure or activation patterns of artificial and biological neural networks. The latter refers to comparing their behavioral outputs (e.g., eye movements, reaction times, high-level decisions). Moreover, comparisons can be made under changing conditions, i.e., during learning and development (Elman et al., 1996). As such, ANNs can serve as explanatory mechanisms in cognitive neuroscience and behavioral psychology, embracing recent model-based approaches (Forstmann and Wagenmakers, 2015).

From a psychological perspective, ANNs have been compared explicitly with their biological counterparts. Connectionist models were widely used in the 1980’s to explain various psychological phenomena, particularly by the parallel distributed processing (PDP) movement, which stressed the parallel nature of neural processing and the distributed nature of neural representations (McClelland, 2003). For example, neural networks have been used to explain grammar acquisition (Elman, 1991), category learning (Kruschke, 1992) and the organisation of the semantic system (Ritter and Kohonen, 1989). More recently, deep neural networks have been used to explain human similarity judgments (Peterson et al., 2016). With new developments in cognitive and affective computing, where neural networks become more adept at solving high-level cognitive tasks, such as predicting people’s (apparent) personality traits (Güçlütürk et al., 2016), their use as a tool to explain psychological phenomena is likely to increase. This will also require embracing insights about how humans solve problems at a cognitive level (Tenenbaum et al., 2011).

ANNs have also been related explicitly to brain function. For example, the perceptron has been used in the modeling of various neuronal systems, including sensorimotor learning in the cerebellum (Marr, 1969) and associative memory in cortex (Gardner, 1988), sparse coding has been used to explain receptive field properties (Olshausen and Field, 1996), topographic maps have been used to explain the formation of cortical maps (Obermayer, 1990; Aflalo, 2006), Hebbian learning has been used to explain neural tuning to face orientation (Leibo et al., 2017), and networks trained by backpropagation have been used to model the response properties of posterior parietal neurons (Zipser and Andersen, 1988). Furthermore, reinforcement learning algorithms used to train neural networks for action selection have strong ties with the brain’s reward system (Schultz et al., 1997; Sutton and Barto, 1998). It has been shown that RNNs trained to solve a variety of cognitive tasks using reinforcement learning replicate various phenomena observed in biological systems (Miconi, 2017; Song et al., 2016). Crucially, these efforts go beyond descriptive approaches in that they may explain *why* the human brain is organized in a certain manner.

Rather than using neural networks to explain certain observed neural or behavioral phenomena, one can also directly fit neural networks to neurobehavioral data. This can be achieved via an indirect approach or via a direct approach. In the *indirect* approach, neural networks are first trained to solve a task of interest. Subsequently, the trained network’s responses are fitted to neurobehavioral data obtained as participants engage in the same task. Using this approach, deep convolutional neural networks trained on object recognition, action recognition and music tagging have been used to explain the functional organization of visual as well as auditory cortex (Güçlü and van Gerven, 2015, 2017; Güçlü et al., 2016). The indirect approach has also been used to train RNNs via reinforcement learning on a probabilistic categorization task. These networks have been used to fit the learning trajectories and behavioral responses of humans engaged in the same task (Bosch et al., 2016). Mante et al. (Mante et al., 2013) used RNNs to model the population dynamics of single neurons in prefrontal cortex during a context-dependent choice task. In the *direct* approach, neural networks are trained to directly predict neural responses. For example, (Mcintosh et al., 2016) trained convolutional neural networks to predict retinal responses to natural scenes, (Joukes et al., 2014) trained RNNs to predict neural responses to motion stimuli, and (Güçlü and Gerven, 2017) used RNNs to predict cortical responses to naturalistic video clips. This ability of neural networks to explain neural recordings is expected to become increasingly important (Sompolinsky, 2014; Marder, 2015), given the emergence of new imaging technology where the activity of thousands of neurons can be measured in parallel (Ahrens et al., 2013; Lopez et al., 2016; Pachitariu et al., 2016; Churchland and Sejnowski, 2016; Yang and Yuste, 2017). Better understanding will also be facilitated by the development of new data analysis techniques to elucidate human brain function (Kass et al., 2014)^10^, the use of ANNs to decode neural representations (Schoenmakers et al., 2013; Güçlütürk et al., 2017), as well as the development of approaches that elucidate the functioning of ANNs (e.g., (Nguyen et al., 2016; Kindermans et al., 2017; Miller, 2017)).^11^

### 5.3 Next-generation artificial neural networks

The previous sections outlined how neural networks can be made to solve challenging tasks and provide explanations of neural and behavioral responses in biological agents. In this final section, we consider some developments that are expected to fuel the next generation of ANNs.

First, a major driving force in neural network research will be theoretical and algorithmic developments that inform why ANNs work so well in practice, what their fundamental limitations are, as well as how to overcome these. From a theoretical point of view, substantial advances have already been made pertaining to, for example, understanding the nature of representations (Anselmi and Poggio, 2014; Lin and Tegmark, 2016), the statistical mechanics of neural networks (Sompolinsky, 1988; Advani et al., 2013), as well as the expressiveness (Pascanu et al., 2013; Bianchini and Scarselli, 2014; Poole et al., 2016; Raghu et al., 2016; Kadmon and Sompolinsky, 2016; Weichwald et al., 2016; Mhaskar et al., 2016) and learnability (Saxe et al., 2014; Dauphin et al., 2014; Schoenholz et al., 2017) of DNNs. From an algorithmic point of view, great strides have been made in improving training of deep (Srivastava et al., 2014; Ioffe and Szegedy, 2015; He et al., 2015) and recurrent neural networks (Hochreiter and Schmidhuber, 1997; Pascanu et al., 2012), as well as on improving the efficacy of reinforcement learning algorithms (Schulman et al., 2015; Mnih et al., 2016; Pritzel et al., 2017).

Second, it is expected that as neural network models become more plausible from a biological point of view, model fit and task performance will further improve (Cox and Dean, 2014). This is important in driving new developments in model-based cognitive neuroscience but also in developing intelligent machines that show human-like behavior. One example is to match the object recognition capabilities of extremely deep neural networks with more biologically plausible RNNs of limited depth (O’Reilly et al., 2013; Liao and Poggio, 2016) and achieving category selectivity in a more realistic manner (Peelen and Downing, 2017; Scholte et al., 2017). Another example is to incorporate predictive coding principles in neural network architectures (Lotter et al., 2016). Furthermore, more human-like perceptual systems can be arrived at by including attentional mechanisms (Mnih et al., 2014) as well as mechanisms for saccade planning (Najemnik and Geisler, 2005; Gregor et al., 2014; Larochelle and Hinton, 2010).

In general, ANN research will benefit from a close interaction between the AI and neuroscience communities (Yuste, 2015). For example, neural network research may be shaped by general guiding principles of brain function at different levels of analysis (O’Reilly, 1998; Maass, 2016; Sterling and Laughlin, 2016). We may also strive to incorporate more biological detail. For example, in order for neural networks to act as accurate models of neural information processing it may be imperative to use spike-based rather than rate-based formulations (Brette, 2015). Efforts are underway to effectively train spiking neural networks (Gerstner et al., 2014; Gerstner and Kistler, 2002; O’Connor and Welling, 2016; Huh and Sejnowski, 2017) and endow them with the same cognitive capabilities as their rate-based cousins (Abbott et al., 2016; Zambrano and Bohte, 2016; Kheradpisheh et al., 2016; Lee et al., 2016; Thalmeier et al., 2015). In the same vein, researchers are exploring how probabilistic computations can be performed in neural networks (Pouget et al., 2013; Nessler et al., 2013; Orhan and Ma, 2016; Heeger, 2017) and deriving new biologically plausible synaptic plasticity rules (Schiess et al., 2016; Brea et al., 2016; Brea and Gerstner, 2016). Biologically-inspired principles may also be incorporated at a more conceptual level. For instance, researchers have shown that neural networks can be protected from adversarial attacks (i.e., the construction of stimuli that cause networks to make mistakes) by incorporating the notion of nonlinear computations encountered in the branched dendritic structures of real neurons (Nayebi and Ganguli, 2016).

Finally, research is invested in implementing ANNs in hardware, also referred to as neuromorphic computing (Mead, 1990). These brain-based parallel chip architectures hold the promise of devices that operate in real time and with very low power consumption (Schuman et al., 2017), driving new advances in cognitive computing (Modha et al., 2011; Neftci et al., 2013). On a related note, nanotechnology may one day drive the development of new neural network architectures whose operation is closer to the molecular machines that mediate the operation of biological neural networks (Drexler, 1992). In the words of Feynman (1992): “There’s plenty of room at the bottom.”

## 6 Conclusion

As cognitive scientists, we live in exciting times. Cognitivism offers an interpretation of agents as information processing systems that reason via physical symbol manipulation. The probabilistic approach to cognition extends this interpretation by viewing organisms as rational agents that need to act in the face of uncertainty under limited resources. Finally, emergentist approaches such as connectionism indicate that concerted interactions between simple processing elements can achieve human-level performance at certain cognitive tasks. While these different views have stirred substantial debate in the past, they need not be irreconcilable. Surely we are capable of formal symbol manipulation and decision making under uncertainty in real-life settings. At the same time, these capabilities must be implemented by the neural circuits that make up our own brains, which themselves rely on noisy long-range communication between neuronal populations.

The thesis of this paper is that natural intelligence can be modeled and understood by constructing artificial agents whose synthetic brains are composed of neural networks. To act as explanations of natural intelligence, these synthetic brains should show a correspondence with their biological counterparts in operation as well as behavior. To identify such similarities we can embrace the rich sources of data provided by neuroscience and psychology. At the same time, we can use sophisticated mathematical machinery developed in machine learning and applied physics to gain a better understanding of these systems. Ultimately, these synthetic brains should be able to show the capabilities that are prescribed by normative theories of intelligent behavior.

The position that artificial neural networks are sufficient for modeling all of cognition may seem exceedingly naive. For example, state-of-the-art question-answering systems such as IBM’s Watson (Ferrucci et al., 2010) use ANN technology as a minor component within a larger largely symbolic framework and the AlphaGo system (Silver et al., 2016), which plays the game of Go at an expert level, combines neural networks with Monte Carlo tree search. While it is true that ANNs remain wanting when it comes to logical reasoning, inferring causal relationships or planning, the pace of current research may very well bring these capabilities within reach in the foreseeable future. Such neural networks may turn out to be quite different from current neural network architectures and their operation may be guided by yet-to-be-discovered learning rules.

The quest for natural intelligence can be contrasted with a pure engineering approach. From an engineering perspective, understanding natural intelligence may be considered irrelevant since the main interest is in building devices that do the job. To quote Edsger Dijkstra, “the question whether machines can think [is] as relevant as the question whether submarines can swim.” At the same time, our quest for natural intelligence will facilitate the development of strong AI given the proven ability of our own brains to generate intelligent behaviour. Hence, biologically inspired architectures can not only provide new insights into human brain function but may also in the long run yield superior curious and perhaps even conscious machines that surpass humans in terms of intelligence, creativity, playfulness and empathy (Moravec, 2000; Modha et al., 2011; Boden, 1998; Der and Martius, 2011; Sze, 2005; Harari, 2017).

## Acknowledgements

This work is supported by a VIDI grant (639.072.513) from the Netherlands Organization for Scientific Research. I would like to thank Nadine Dijkstra, Andrew Reid and Katja Seeliger for their useful comments on a previous version of this manuscript.

## Footnotes

1 In fact, ACT-R also uses some subsymbolic elements and can therefore be regarded a *hybrid* architecture.

2 Beliefs over continuous quantities can be expressed by replacing summation with integration.

3 From: https://www.izhikevich.org/human_brain_simulation/why.htm

4 Intentionality or “aboutness” referes to the quality of mental states as being directed towards an object or state of affairs.

5 In practice, it is more efficient to iterate over subsets of datapoints, known as mini-batches, in sequence. That is, training is organized in terms of epochs in which all datapoints are processed by iterating over mini-batches. Note that, whenever we are not processing all data points in parallel, we are not exactly following the gradient. Therefore, any such procedure is known as *stochastic gradient descent*.

6 The ability of simple RNNs to integrate information over time remains limited, which led to the introduction of various extensions that perform more favorably in this regard (Hochreiter and Schmidhuber, 1997; Cho et al., 2014; Wu et al., 2016; Neil et al., 2016).

7 As formulated by Lake and Tenenbaum (Lake et al., 2016): “The difference between pattern recognition and model-building, between prediction and explanation, is central to our view of human intelligence. Just as scientists seek to explain nature, not simply predict it, we see human intelligence as a model-building activity.”

8 See Winograd’s SHRDLU for an early example of such a virtual world (Winograd, 1972).

9 The notion of *wanting* agents was already present in the writings of Thurstone, who wrote the following (Thurstone, 1923): “My main thesis is that conduct originates in the organism itself and not in the environment in the form of a stimulus. […] All mental life may be looked upon as incomplete behavior which is in the process of being formed. […] Perception is the discovery of the suitable stimulus which is often anticipated imaginally. The appearance of the stimulus is one of the last events in the expression of impulses in conduct. The stimulus is not the starting point for behavior.”

10 But see (Jonas and Kording, 2017) for a critical appraisal of the informativeness of such techniques.

11 These techniques aim to overcome the interpretability problem raised by Mozer and Smolensky (1989), who state: “One thing that connectionist networks have in common with brains is that if you open them up and peer inside, all you can see is a big pile of goo.”

